# ERK signalling orchestrates metachronous transition from naïve to formative pluripotency

**DOI:** 10.1101/2023.07.20.549835

**Authors:** Carla Mulas, Melanie Stammers, Siiri I. Salomaa, Constanze Heinzen, David M. Suter, Austin Smith, Kevin J. Chalut

## Abstract

Naïve epiblast cells in the embryo and pluripotent stem cells *in vitro* undergo developmental progression to a formative state competent for lineage specification. During this transition, transcription factors and chromatin are rewired to encode new functional features. Here, we examine the role of mitogen-activated protein kinase (ERK1/2) signalling in pluripotent state transition. We show that a primary consequence of ERK activation in mouse embryonic stem cells is elimination of Nanog, precipitating breakdown of the naïve state gene regulatory network. Cell variability in pERK dynamics results in metachronous down-regulation of Nanog and naïve state exit. Knockdown of Nanog allows exit without ERK activation. However, transition to formative pluripotency does not proceed and cells collapse to an indeterminate identity. This failure is attributable to loss of expression of the central pluripotency factor Oct4. Thus, during formative transition ERK signalling both dismantles the naïve state and preserves pluripotency. These results illustrate that a single signalling pathway can both drive exit from a developmental state and safeguard progression to the successor state.

## INTRODUCTION

How cells traverse between proximate yet molecularly and functionally distinct identities is a long-standing question in developmental and stem cell biology. Pluripotent mouse embryonic stem cells (ESCs) reside in a naïve state. To undergo germline and somatic differentiation, they must first transition to a formative state that is competent to respond to lineage induction (Hayashi et al., 2011; Kinoshita et al., 2021; Mulas et al., 2017). In the naïve state, ESCs undergo continuous symmetrical self-renewal while retaining pre-implantation epiblast identity and the ability to colonise blastocyst chimaeras (Bradley et al., 1984). Stable propagation of naïve ESCs is achieved using “2i” culture conditions, comprising small molecule inhibition of MEK upstream of ERK1 and ERK2 (henceforth ERK) and partial inhibition of glycogen synthase kinase 3 (GSK3) (Ying et al., 2008) Leukaemia inhibitory factor (LIF) further increases self-renewal efficiency. Withdrawal of 2i and culturing cells in only N2B27 basal media, opens the path to differentiation via transition to formative pluripotency (Smith, 2017). Formative pluripotent cells have distinct molecular, cell biological and developmental properties compared to naïve cells. They have different transcription factor dependencies, gene expression signatures, chromatin modifications, DNA methylation, metabolism and cell surface mechanics (de Belly et al., 2021; Hayashi et al., 2011; Kalkan et al., 2017a; Kinoshita et al., 2021; Mulas et al., 2017; Smith, 2017; Wang et al., 2021; Yu et al., 2021). Most importantly, while naïve cells cannot respond productively to lineage-inducing signals, formative cells have gained the competence to undergo efficient germline and germ layer lineage specification (Hayashi et al., 2011; Kinoshita et al., 2021; Mulas et al., 2017). Similar findings have been confirmed for human naïve pluripotent stem cells (Rostovskaya et al., 2019).

Multiple studies have established that the ERK pathway acting downstream of autocrine FGF4 is a key driver of the pluripotency transition. Perturbations in components of the FGF/ERK pathway prevent or significantly impede differentiation (de Belly et al., 2021; Betschinger et al., 2013; Burdon et al., 1999; Cheng et al., 1998; Findlay et al., 2013; Kunath et al., 2007; Leeb et al., 2014; Molotkov et al., 2017; Sangokoya and Blelloch, 2020; Stavridis et al., 2007; Yang et al., 2012). Inhibition of this pathway thus empowers the robust derivation and propagation of ESCs (Batlle-Morera et al., 2008; Ying et al., 2008) Conversely, increasing ERK activity by modulating negative feedback regulators, such as the ribosomal S6 kinase (RSK) family, leads to a more rapid and synchronised progression from naïve to formative cell states (Nett et al., 2018; Yang et al., 2012). A recent study using optogenetic stimulation of FGFR1 has further confirmed that ERK activity propels naïve state exit (Arekatla et al., 2023). How this is achieved remains unclear, however.

Here, we combine chemical modulation of ERK activity with transcription factor perturbations to examine the requirements for both naïve state exit and progression to formative pluripotency.

## RESULTS

### Kinetics of naïve state exit depends on activation of ERK

To monitor the effect of ERK activity on exit from the naïve ESC state, we employed two assay systems: (i) colony formation on replating in 2iLIF (Dunn et al., 2019); (ii) expression of the *Rex1:GFPd2* (RGd2) fluorescent reporter, specific for the naïve state (Kalkan et al., 2017a). We initially validated these assays by examining cells withdrawn from 2i for 30h, culturing cells in N2B27 basal media. In agreement with previous results (Nett et al., 2018), treatment with the MEK inhibitor [MEK(i)] PD0325901 to block ERK activation was sufficient to fully maintain ESC colony formation ability after 30h (Figure S1A). Cells also retained expression of the naïve RGd2 reporter in MEK(i) in similar proportion to cells maintained in 2i (Figure S1B). To examine the effect of increased ERK signalling, we blocked negative regulation by RSK proteins using the pan-RSK inhibitor [RSK(i)] BI-D1870 (Nett et al., 2018). Following treatment with RSK(i) for 30h, cells formed fewer ESC colonies (Figure S1A) and showed more extensive downregulation of the RGd2 reporter than cells in medium only (Figure S1B). Thus, we confirmed that ERK1/2 signalling activity regulates the rate of exit from the naïve ESC state (Nett et al., 2018).

Mouse ESCs are sustained by cooperative and partially redundant activity of a suite of transcription factors that act in conjunction with the constitutive pluripotency factors Oct4 and Sox2 (Dunn et al., 2014). Among these, Klf4 and Nanog have been reported to be directly phosphorylated and potentially destabilised by pERK (Dhaliwal et al., 2018; Kim et al., 2014). We used siRNA to assess whether depletion of either factor, or of Esrrb which lies downstream of Nanog (Festuccia et al., 2012), would allow naïve state exit without ERK signal (Figure 1A). *siKlf4* showed no effect on RGd2 expression at 30h in the presence of MEK(i). In contrast, treatment with *siNanog* or *siEsrrb* resulted in RGd2 downregulation (Figure 1B, S1C) and furthermore reduced colony formation on replating (Figure 1C). Therefore, the ability of MEK(i) to delay naïve state exit requires *Nanog* or *Esrrb*. We also examined the consequences of knockdown on transition when ERK signalling is active (Figure 1D). Both *siNanog* and *siEsrrb* resulted in faster downregulation of RGd2 compared to scramble control siRNA or *siKlf4* (Figure 1E). These data indicate that clearance of *Nanog* and *Esrrb* is rate-limiting for exit from the naïve ESC-state.

**Figure 1.**
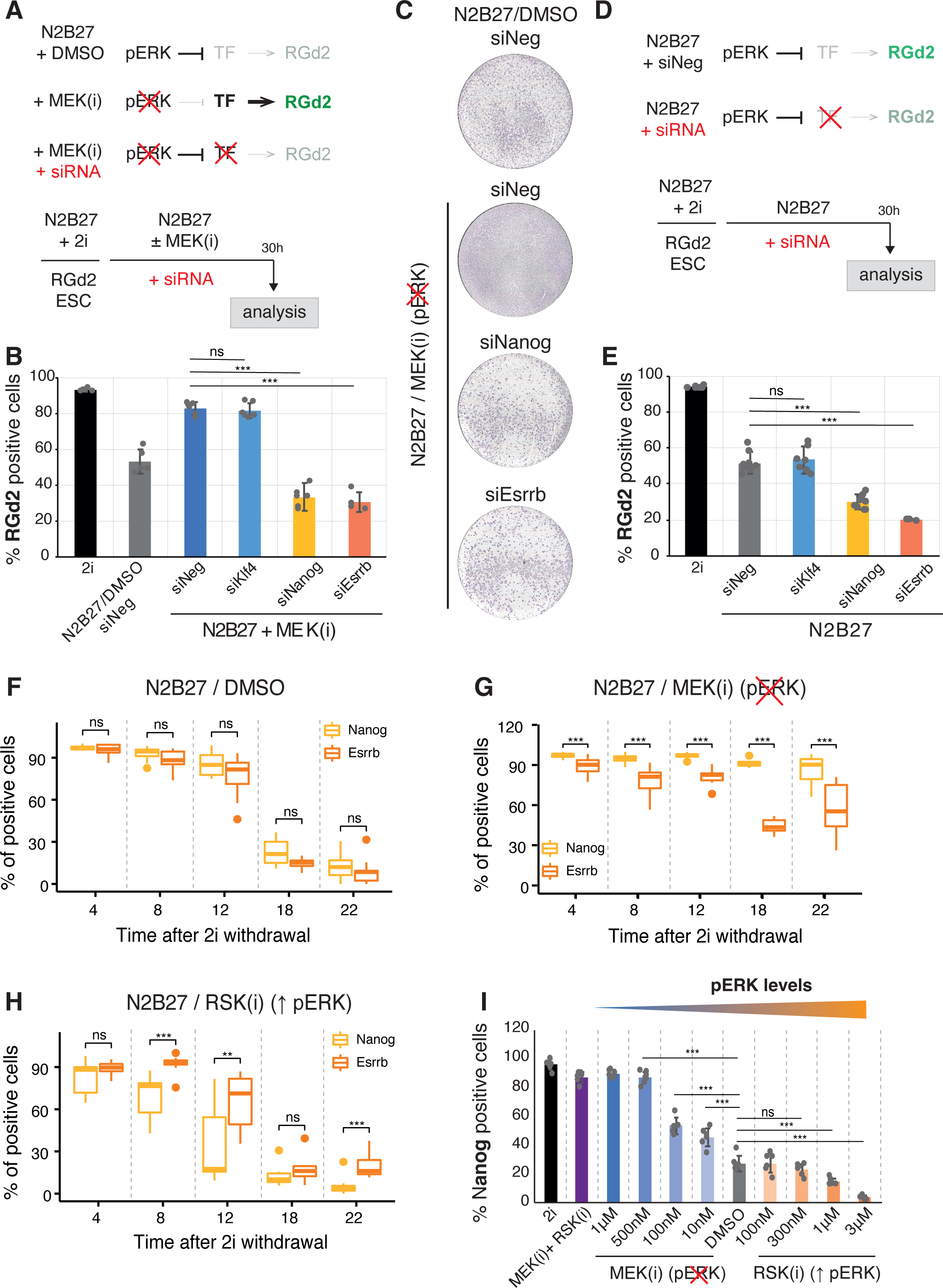
Identifying transcription factors downstream of ERK. (A) Strategy to identify transcription factors downstream of ERK. (B) Percentage of RGd2 positive cells determined by flow cytometry, 30 hours after 2i withdrawal. Mean ±SD, student t-test with respect of N2B27/MEK(i)+siNeg (N=6). (C) Representative images of colony formation in 2i/LIF conditions of 30-hour cells. (D) Strategy to identify rate-limiting steps during exit from naïve pluripotency. (E) Percentage of RGd2 positive cells determined by flow cytometry following siRNA treatment in N2B27 conditions. Mean ±SD, student t-test with respect of N2B27/MEK(i)+siNeg (N=6). (F-H) Boxplots showing percentage of Nanog and Esrrb positive cells over time in N2B27/DMSO under endogenous ERK activation (F), in N2B27/MEK(i) (G) and in N2B27/RSK(i) conditions (H) (N=2, 6 images/condition). Two-sided Wilcoxon test. (I) Percentage of Nanog positive cells at 20 hours under different concentrations of MEK(i) and RSK(i). (N=2, 6 images/condition). Student t-test. Statistics: ns (not significant), * p < 0.05, ** p < 0.005, *** p < 0.0005.

We examined the effect of ERK perturbations on Nanog and Esrrb protein by immunofluorescence staining. In medium alone downregulation of both proteins occurred between 12h and 18h (Figure 1F, S1D). In MEK(i), nearly all cells remained strongly positive for Nanog protein for at least 22h (Figure S1E) whereas Esrrb was diminished in around 50% of cells by 18h (Figure 1G). Conversely, enhancing ERK activity with RSK(i) treatment resulted in earlier downregulation of Nanog than Esrrb (Figure 1H). Therefore, ERK activity elicits loss of Nanog protein before Esrrb.

To determine if Nanog responsiveness is dose-dependent, we titrated MEK(i) and RSK(i) and measured the percentage of Nanog positive cells at 20hrs. We found that higher levels of ERK activity resulted in a lower number of Nanog positive cells (Figure 1I). Treatment with both MEK(i) and RSK(i) phenocopied MEK(i)-only, indicating that the effect of RSK(i) on Nanog is through the ERK pathway (Figure 1I).

We asked how the time of ERK activation correlates with Nanog down-regulation in individual cells since both processes are heterogenous in the population (Arekatla et al., 2023; Reimann et al., 2023). To capture the dynamics of ERK activity over the timescale of cell state transitions, we monitored expression of a direct transcriptional target, *Spry4*. We employed ultrasensitive bioluminescence live imaging, which has low phototoxicity, and used luciferase reporters with fast folding and degradation kinetics (Mandic et al., 2017). Using ESCs carrying a *Spry4-Fluc* gene trap and a previously validated *Nanog::Nlu*c protein fusion knock-in, we imaged at 10 min intervals. Withdrawal of 2i/LIF resulted in dynamic upregulation of the Spry4-Fluc reporter, consistent with dynamic ERK activation profiles observed in ESCs and other cells (Figure 2A, B) (Albeck et al., 2013; Aoki et al., 2013; Nett et al., 2018). Of interest, we observed only one Spry4-Fluc peak in most cell division cycles.

**Figure 2.**
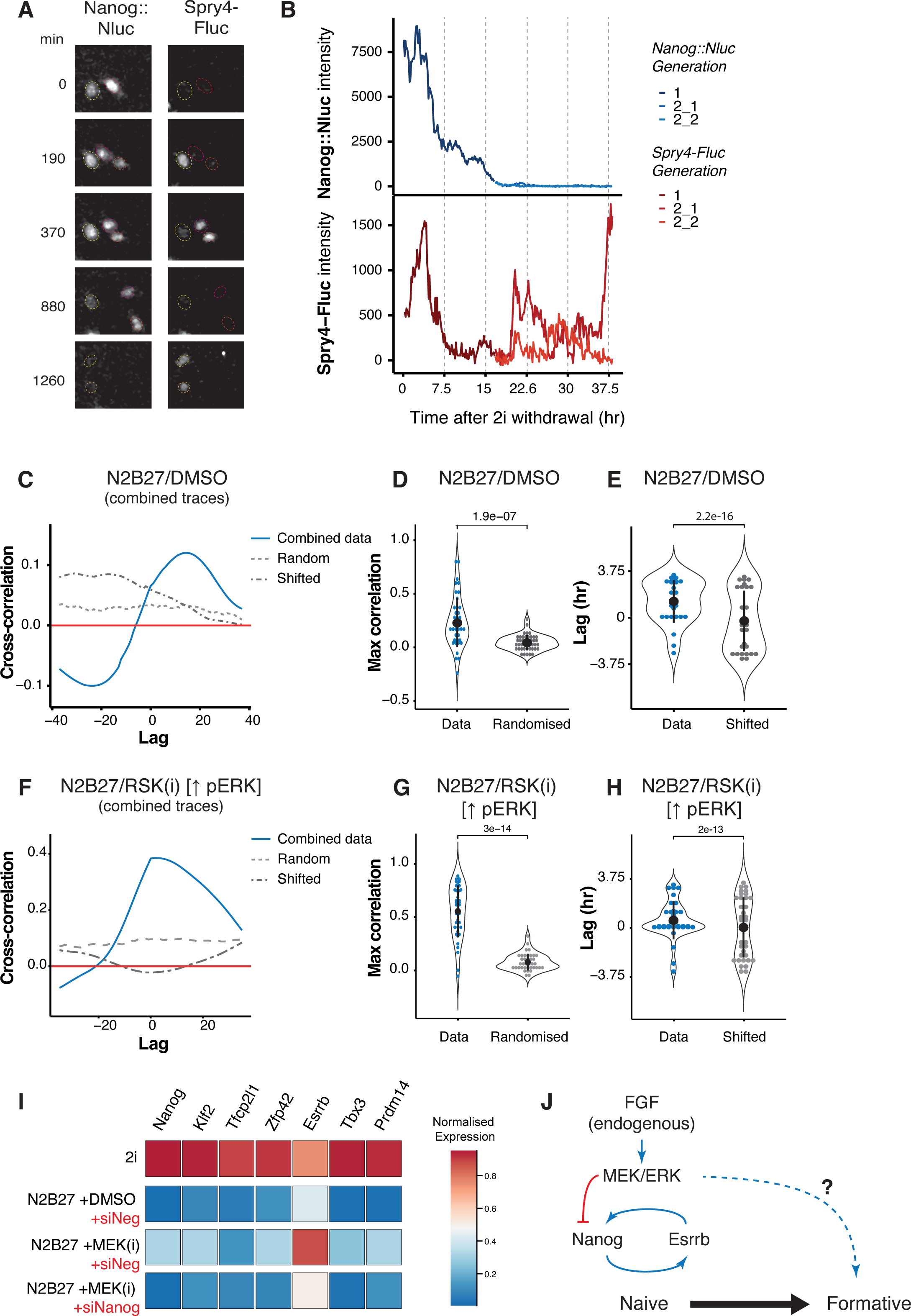
Live imaging ERK activity and Nanog protein dynamics. (A) Representative image of Spry4-Fluc, Nanog::Nluc signal after withdrawal of 2i culture conditions. Red, orange and Yellow circles show different cells across selected frames. (B) Representative trace of Nanog::Nluc and Spry4-Fluc over ∼38hours of differentiation. (C) Autocorrelation function plot between Spry4 promoter activity and Nanog protein for the combined data (all traces joined end-to-end), as well as the randomised and shifted controls in N2B27/DMSO conditions. (D) Comparison of maximum ACF between Spry4 promoter and Nanog protein in individual traces against the randomised control in N2B27/DMSO conditions. In black is mean ± SD. Student t-test (n=44, two independent imaging experiments). (E) Comparison of peak Lag time at maximum ACF for all the cells that show a significant max ACF above noise compared to traces where the Spry4 promoter activity was randomly shifted. In black mean ± SD. Statistics show comparison of the distributions using Kolmogorov– Smirnov (n=28, two independent imaging experiments). (F) Autocorrelation function plot between Spry4 promoter activity and Nanog protein for the combined data (all traces joined end-to-end), as well as the randomised and shifted controls in N2B27/RSK(i) conditions. (G) Comparison of maximum ACF between Spry4 promoter and Nanog protein in individual traces against the randomised control in N2B27/RSK(i) conditions. In black is mean ± SD. Student t-test (n=38, two independent imaging experiments). (H) Comparison of peak Lag time at maximum ACF for all the cells that show a significant max ACF above noise compared to traces where the Spry4 promoter activity was randomly shifted in N2B27/RSK(i) conditions. In black mean ± SD. Statistics show comparison of the distributions using Kolmogorov–Smirnov (n=35, two independent imaging experiments). (I) Expression of naïve genes by RT-qPCR of 30hr cells under different conditions. Expression was normalised to *Actb*, then to the maximum for each gene. (N=3). (J) Schematic of the proposed link between ERK and the Nanog/Esrrb axis in maintaining naïve pluripotency. Statistics: ns (not significant), * p < 0.05, ** p < 0.005, *** p < 0.0005.

To study the relationship between Spry4 activation and Nanog downregulation, we filtered the dataset to exclude timepoints after Nanog is downregulated. After smoothing to remove noise, we used a simple set of ordinary differential equations to calculate the *Spry4* promoter activity for each Spry4-Fluc trace (Figure S2A, see methods for details). We then examined the correlation between the *Spry4* promoter activity and the Nanog:Nluc signal by calculating the cross-correlation (Figure S2B, C). We observed a strong correlation between the signals compared to randomised data in the combined dataset (Figure 2C, S2B) and for individual traces (Figure 2D, S2D). For all cells that showed significant correlation between Spry4 promoter activity and Nanog, we also observed a more consistent and significant difference in the time offset or lag between Nanog downregulation and Spry4 activation compared to randomly shifted controls (Figure 2E, S2E). The average lag time is ∼80min, indicating that activation of the *Spry4* promoter precedes Nanog downregulation. We repeated the analysis for RSK(i) treated cells and observed a stronger correlation at the level of the combined dataset (Figure 2F) as well as in individual cells (Figure 2G, S2F). Interestingly, RSK(i) treatment decreased the average lag between *Spry4* promoter activation and Nanog downregulation to 20 min (Figure 2H, S2G). Since the lag is very short, and not evident in all cells, Nanog downregulation might not require prior transcriptional activation.

Overall, these analyses indicate that the level or dynamics of pERK that activate a transcriptional response in individual cells also initiate downregulation of Nanog protein. Thus, metachronous exit from the naïve state is determined by cell-to-cell variability in pERK activation.

We analysed mRNA level of key naive ESC markers by RT-qPCR (Figure 2I). These markers are expressed highly in 2i and downregulated after 30h withdrawal. Cells maintained in MEK(i) showed partial reduction, ∼70%, for Nanog and other factors, except for *Esrrb* which surprisingly exhibited increased mRNA (Figure 2I). These findings contrast with the protein dynamics we observed above (Figure 1G). *Esrrb* is a well-characterised direct transcriptional target of Nanog (Festuccia et al., 2012). We therefore combined MEK(i) with si*Nanog*. In that condition we observed loss of *Esrrb* mRNA by 30h (Figure 2I), consistent with the observed naïve state exit.

These findings reveal that during pluripotency transition the proximal effect of acute ERK activation on the naïve transcription factor network is to reduce Nanog protein, which in turn diminishes *Esrrb* transcription. This sequence of events is predicted to collapse the naïve gene regulatory network and propel exit from the naïve state (Figure 2J) (Dunn et al., 2014; Festuccia et al., 2018).

### Ongoing ERK activation is required for entry into formative pluripotency

On release from 2i, ERK signalling downregulates the Nanog/Esrrb axis and allows cells to exit the naïve state (Figure 3A). When MEK(i) is maintained to block ERK activation, cells do not upregulate formative genes by 30h, consistent with persistence in the naïve state (Figure 3B, D). However, depletion of Nanog allows us to enforce naïve state exit in the absence of ERK activity. We investigated whether exit in these conditions is sufficient for transition to the formative state (Figure 3C). We found that upon Nanog knockdown and collapse of the naïve transcription factor network, cells maintained in MEK(i) remained viable but failed to upregulate formative genes, notably including *Otx2* a key effector of formative pluripotency (Figure 3D) (Acampora et al., 2012; Buecker et al., 2014). This indicates an ongoing requirement for ERK signalling to execute the transition and establish formative pluripotency.

**Figure 3.**
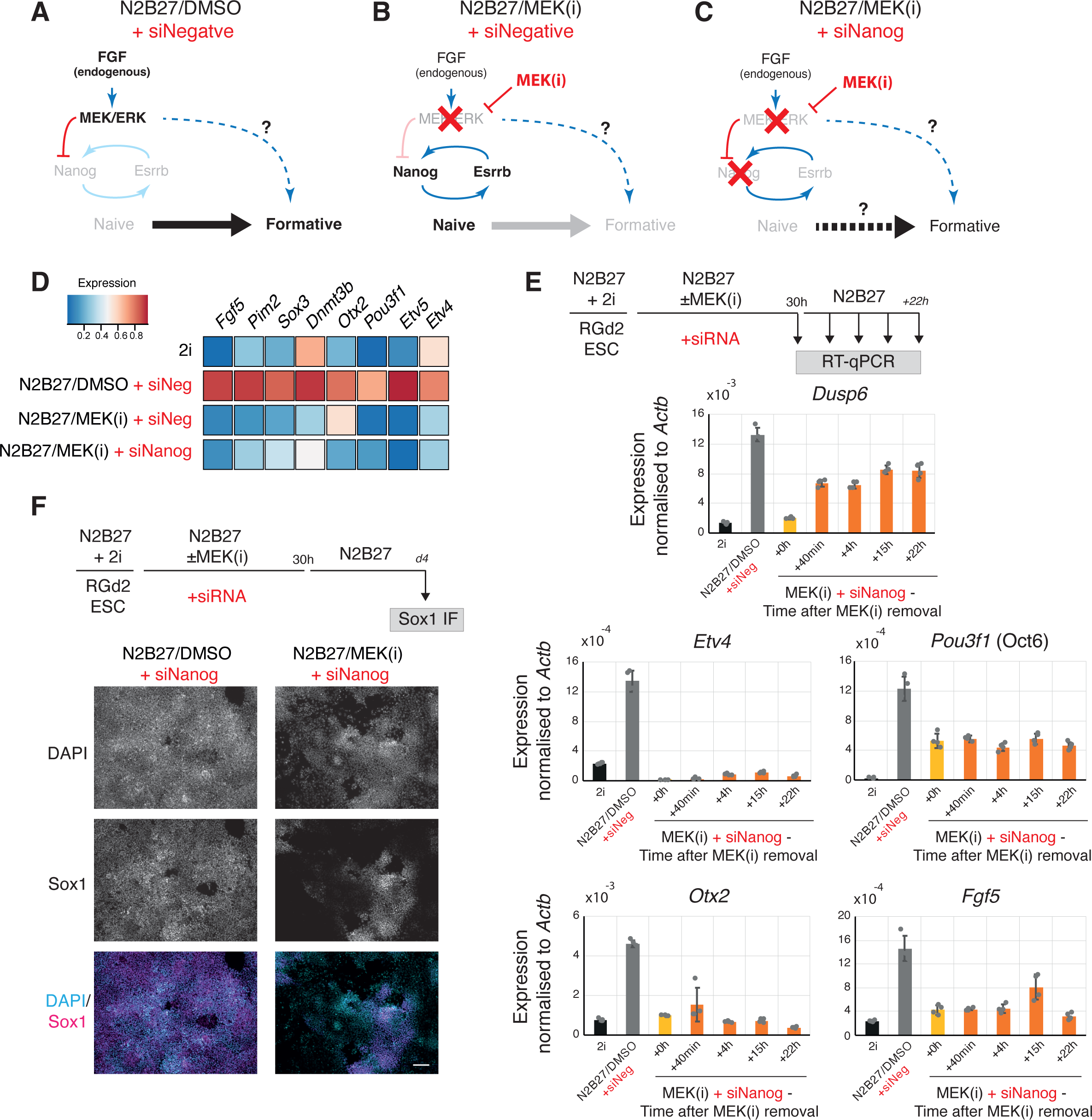
ERK is required for entry into formative state. (A-C) Schematics summarising the effect of the key culture conditions. (D) Expression of formative genes by RT-qPCR under different conditions at 30hrs. Expression was normalised to *Actb*, then to the maximum for each gene. Heatmap of means (N=3). (E) Experimental set up and results to rescue expression of formative genes following withdrawal of MEK(i). Expression normalised to Actb. Mean ± SD (N=4). (F) Representative immunostaining for Sox1 expression on day 4 of neural differentiation of siNanog cells treated with DMSO or MEK(i). Scale bar = 50 μm

We investigated whether MEK(i)+siNanog cells may be stalled in transition and poised for formative progression upon ERK activation. We therefore removed MEK(i) after 30h to release ERK signalling. However, we saw no increase in expression of *Otx2* or other formative marker genes (Figure 3E). We also assessed whether N2B27/MEK(i)+siNanog cells may be diverted into an extraembryonic fate. However, we did not detect significant expression of trophoblast markers *Cdx2*, *Eomes* or *Hand1* (Figure S3A). Similarly, we could not detect appreciable expression of endoderm markers *Sox17*, *Gata6* or *Gata4*, ruling out both hypoblast and definitive endoderm differentiation (Figure S3B).

We considered whether MEK(i)+siNanog cells might experience precocious differentiation, truncating or bypassing the formative stage. To test this possibility, siNanog cells were treated with MEK(i) before withdrawing the inhibitor and measuring expression of Sox1, a neural specific transcription factor, on day 4. Culture of siNanog cells without MEK(i) resulted in efficient upregulation of Sox1 on day 4 (Figure 3F). This confirms that cells exiting naïve pluripotency without MEK(i) can gain competence for lineage specification and that Nanog is not required for this. However, treatment of siNanog cells with MEK(i) for 30hrs substantially impaired upregulation of Sox1 (Figure 3F). Therefore, cells do not undergo accelerated neural differentiation after naïve state exit in the presence of MEK(i). By removing MEK(i) at different timepoints, we observed that treatment for longer than 18hrs was sufficient to significantly impair neural differentiation (Figure S3C).

In summary, MEK(i)+siNanog cells downregulate ESC markers and exit the naïve state, but they neither upregulate formative genes nor divert to extraembryonic identity. These findings indicate that ERK signalling is necessary after exit for progression to the formative state and that without ERK input cells become stranded in an indeterminate state.

### Failure to transition is not due to genome-wide chromatin dysregulation

ERK signalling has been shown to have effects on chromatin and transcription (Tee et al., 2014). Therefore, we asked whether MEK(i)+siNanog cells were disabled by a global alteration of the chromatin landscape. Using Cut&Tag (Kaya-Okur et al., 2019) we examined the distribution of H3K27me3, H3K4me3 and H3K27ac histone modifications. During normal transition, most H3K27me3 promoter peaks are maintained at 30hrs, and only 7-10% differ (lost or acquired) (Figure S4A). In MEK(i)+siNanog the vast majority of H3K27me3 promoter peaks are similarly maintained. Therefore, the failure in formative transition is unlikely to be due to global dysregulation of the repressive chromatin landscape. However, approaching 50% of the differential sites were alternatively modulated (Figure S4B, S4C). Amongst promoter regions showing an aberrant increase in H3K27me3, many are associated with ERK responsive genes such as *Dusp6* or with formative genes like *Pou3f1* (Figure S4F). A small proportion of promoters, including *Myc,* gained H3K27me3 only in MEK(i)+siNanog conditions, (Figure S4C). H3K27me3 redistribution at enhancers was similar to that observed at promoters, with most peaks unchanged in either condition, and of those that do change, only a small proportion behaving differently in MEK(i)+siNanog (Figure S4D).

H3K27ac deposition is more dynamic during transition. H3K27ac peaks present in 2i were lost at more than half of sites with only a quarter maintained and ∼20% gained (Figure S4E). This pattern was not dramatically different in MEK(i)+siNanog treated cells, although H3K27ac was not gained at ∼30% of appropriate sites (Figure S4E), including the ERK target and formative marker *Lef1* (Figure S4F).

Overall, these data do not indicate global dysregulation of histone modifications due to ERK pathway inhibition. Instead, the data show that many ERK response genes and formative genes acquire increased H3K27me3 and lower H3K4me3 and H3K27ac (Figure S4F), while a smaller group of genes fail to gain H3K27me3 and lose H3K4me3 and H3K27ac (Figure S4G). These differences are consistent with, and potentially consequent to, reduced transcription of those genes. However, higher H3K27me3 deposition at promoters and enhancers of many formative genes may contribute to the inability to restore expression by removing MEK(i) after 30h (Figure 3E).

### ERK activity is required to maintain Oct4 expression in the naïve to formative pluripotency transition

The transcription factor Oct4 (gene name *Pou5f1*) is the mainstay of pluripotency, expressed through all stages *in vivo* and *in vitro* (Schöler et al., 1990; Wu and Schöler, 2014). In the preceding analyses of histone modifications, we noticed specific changes at the Oct4 promoter, which prompted further investigation. In MEK(i)+siNanog cells, we detected loss of H3K4me3 and to a lesser extent of H3K27ac at the *Pou5f1* locus (Figure 4A). Oct4 expression is normally maintained throughout the naïve to formative transition. However, in line with disappearance of H3K4me3, we observed that MEK(i)+siNanog cells did not express Oct4 mRNA (Figure 4B) and also lost Oct4 protein (Figure 4C, D). Furthermore, release from MEK(i) did not reactivate Oct4 expression (Figure 4B). We similarly saw that MEK(i)+siEsrrb cells failed to maintain Oct4 protein (Figure S5A). To determine when cells lose Oct4 expression, we measured the percentage of Oct4 positive cells over time in N2B27/DMSO+siNeg and in N2B27/MEK(i)+siNanog. In N2B27, Oct4 protein diminishes between ∼36-48hrs after 2i withdrawal, as cells begin to upregulate Sox1 and enter the neural lineage (Figure 4C) (Mulas et al., 2017). In contrast, in MEK(i)+siNanog cells Oct4 negative cells are detected as early as 18h after 2i-withdrawal and expression is extinguished throughout the population by 30h (Figure 4C, D).

**Figure 4.**
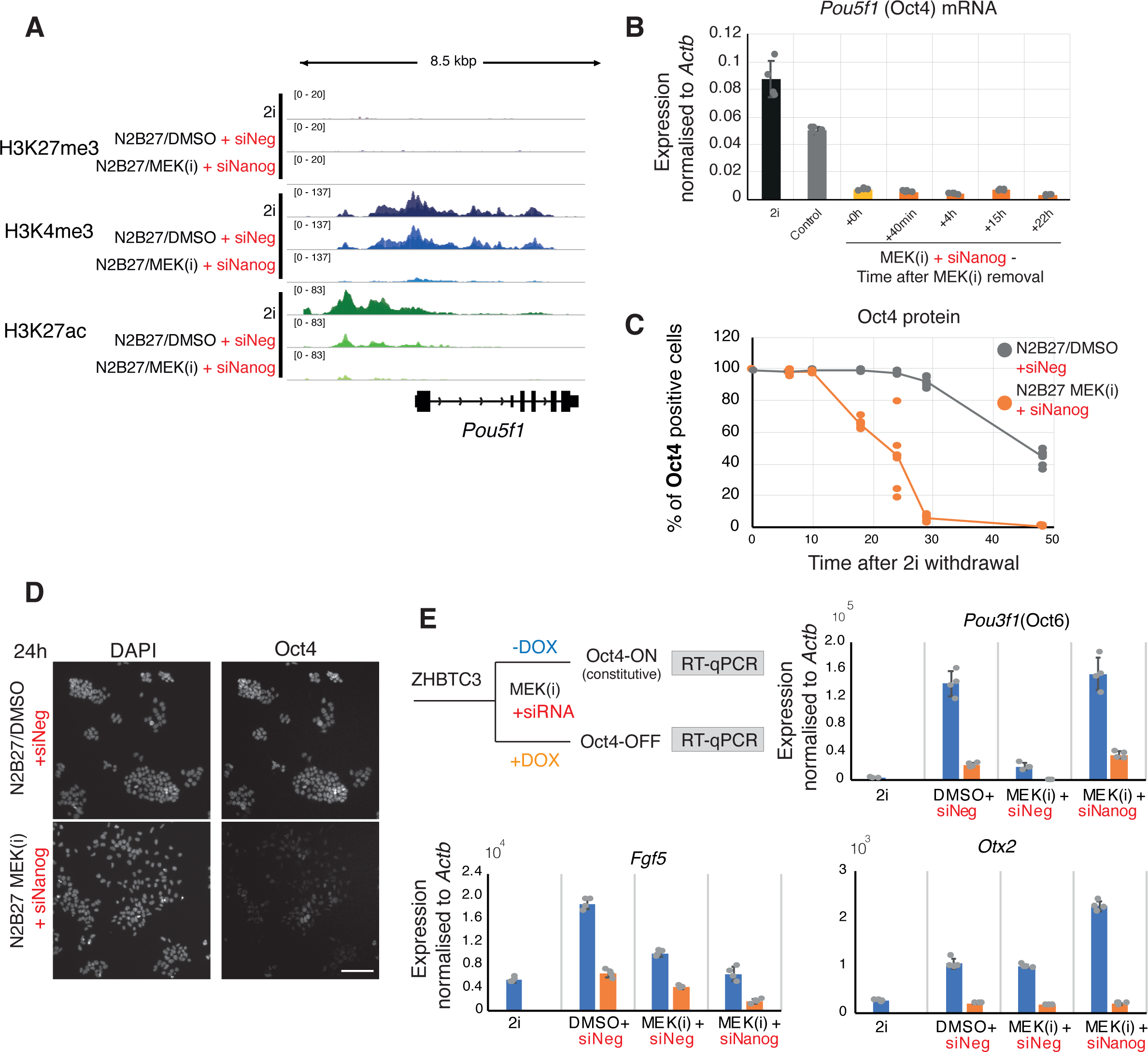
Failure to maintain Oct4 expression contributes to defective entry into the formative state. (A) Distribution of H3K27me3, H3K4me3 and H3K27ac along the Oct4 locus. (B) mRNA levels determined by RT-qPCR of Pou5f1 (Oct4) in control, N2B27/MEK(i) + siNanog cells and after withdrawal of MEK(i). Mean ±SD shown (N=4). (C) Percentage of Oct4 positive cells as determined by immunofluorescence over time in N2B27/DMSO and N2B27/MEK(i) + siNanog conditions. (D) Representative image showing immunofluorescence for Oct4 protein in N2B27/DMSO + siNegative and N2B27/MEK(i) + siNanog conditions at 24hrs. Error bar: 100μm. (E) Rescue of expression of formative genes by constitutive expression of Oct4. Experimental set up (top left) and results. Graphs show gene expression determined by RT-qPCR, normalised to *Actb* (N=4).

We investigated whether constant expression of Oct4 is sufficient to restore entry into the formative state in MEK(i)+siNanog conditions. For this we used ZHBTc4 ESCs, in which both Oct4 alleles are inactivated and Oct4 is produced from a regulatable transgene (Niwa et al., 2000). Expression of the transgene is repressed by addition of doxycycline (DOX) (Figure 4E). Without DOX (Oct4 ON), ZHBTc4 cells efficiently upregulate formative genes after withdrawal from 2i (Figure 4E). If DOX is added to silence Oct4 expression, ZHBTc4 cells become trophectoderm-like (Niwa et al., 2000). In this condition formative genes are not activated, confirming that Oct4 is necessary for formative transition. We then treated ZHBTc4 cells with MEK(i)+siNanog in either the absence or presence of DOX. When DOX was added to silence Oct4, cells failed to upregulate formative genes as with control-treated cells. In contrast, without DOX (Oct4 ON) MEK(i)+siNanog ZHBTc4 cells displayed upregulation of key formative transcription factors *Otx2* and *Pou3f1*. An exception that did not show restored expression is *Fgf5*, likely because it is a direct target of ERK (Kalkan et al., 2017b).

We conclude that ERK1/2 activity becomes essential for continued Oct4 expression as cells exit the naïve state, and that without Oct4 cells are unable to upregulate formative transcription factors and proceed with transition.

Collectively these results demonstrate that ERK1/2 signalling drives ESC transition both by destabilising the naïve state transcription factor network and by sustaining the platform pluripotency factor Oct4. The detailed mechanisms of these two distinct effects remain to be determined. It is known that pERK can phosphorylate Nanog (Brumbaugh et al., 2014; Kim et al., 2014) but whether this is sufficient to acutely destabilise the protein has yet to be determined. How pERK regulates Oct4 gene expression requires further investigation, but we speculate that this may be linked to the enhancer switching that is known to occur at the Oct4 locus during formative transition (Choi et al., 2016; Tesar et al., 2007; Yeom et al., 1996).

In summary, our findings show that variable dynamics of ERK activation in ESCs trigger metachronous dissolution of the naïve state and that continuation of ERK signalling preserves expression of the anchor factor(s) Oct4 to secure cell state transition.

## METHODS

### Culture of mouse embryonic stem cells

Mouse embryonic stem cells were routinely cultured in 2i (1 μM PD0325901 [MEK(i)], 3 μM CHIR99021) or 2i with LIF (1:1000 LIF Qkine) following the protocols outlined in (Mulas et al., 2019). Cells were dissociated using Accutase (Millipore, SCR005) and pipetted to obtain a single cell suspension. The cell suspension was diluted in 5x wash buffer [DMEM/F12, 0.03% BSA Fraction V (Thermo Fisher)], span down and resuspended in fresh 2i or 2i/LIF medium before cell counting. Cells were plated at 1.5x10^4^ cells/cm^2^. For maintenance, cells were plated on dishes coated with 0.1% gelatine for at least 15min at 37°C (Sigma-Aldrich, G1890), while all exit and differentiation experiments were performed in plates coated with 10μg/ml laminin in PBS for at least 30 min at 37°C (Sigma Millipore CC095). For all experiments, batch tested, home-made N2B27 was used [N2B27.BV, (Mulas et al., 2019). Unless indicated, the following inhibitors were used: MEK(i) PD0325901, 1 μM; RSK(i) BI-D1870 (Axon Medchem, Axon 1528), 3 μM; For experiments with the Zhbtc2 line, 1μg/ml doxycycline (Sigma) was added to the media when indicated.

### siRNA treatment and exit from naïve state

All siRNA experiments were performed in 24 well plates in technical duplicates and biological triplicates, following the protocols indicated in (Mulas et al., 2019). For each well, we plated 30,000 cells, 20nM of total siRNA and 0.5μl Lipofectamine RNAiMax (Thermo Fisher) in 500μl 2i. After overnight incubation (<18hrs), cells were either collected for RNA extraction and cDNA synthesis to determine the efficiency of knockdown or further differentiated. For differentiation experiments, cells were gently washed with 1ml PBS, before adding fresh media (e.g. N2B27 ± inhibitors). For each gene we used 2 siRNAs (10nM each). Samples were analysed at the indicated timepoints. The list of siRNAs used is in Table S1, and were previously validated in (Dunn et al., 2014).

### RT-qPCR

RNA was extracted using Qiagen RNeasy or Relia Prep RNA Miniprep System (Promega). cDNA synthesis was performed using SuperScript III (Invitrogen) following manufacturers protocols. Quantitative PCR (qPCR) was performed using TaqMan (Thermo Scientific), Sybr Green (Thermo Scientific) or UPL (Roche) technology. A list of primers and probes is provided in Table S1.

### Immunostaining and quantification

Samples were fixed with cold 4% PFA (Santa Cruz Biotechnology) at room temperature for 10 min and washed twice with PBS. Samples were stored in PBS at 4°C until all timepoints could be stained in parallel. Samples were permeabilised and blocked in block buffer (0.3% donkey serum, 0.15% Triton X in PBS) for at least 2 hours at room temperature. Primary antibodies were diluted in block buffer and incubated overnight at 4°C (Table S1). Samples were washed three times with PBST (0.15% Triton X in PBS) for 15min each time. Alexa conjugated secondary antibodies (Invitrogen) were diluted 1:1000 in block buffer alongside a nuclear stain (DAPI or Hoechst) and incubated for 2 hours at room temperature. Samples were washed three times in PBST for 15min each before storing samples overnight in PBS at 4°C.

Images of fixed samples were acquired using a Leica DMI4000 at 10x or 20x magnification. For each biological repeat of a time series, all images were acquired in a single imaging session. For each well, we acquired 3-8 images in a grid pattern. Images processing and segmentation was performed in CellProfiler (https://github.com/CMulas/IF_quantification). Nuclear stain was used to segment nuclei before determining the intensity on the remaining channels. The intensity measures were then analysed using R code (https://github.com/CMulas/IF_quantification). We used receiver operating characteristic (ROC) curve to determine a binary threshold between positive and negative control cells. Whenever possible, negative control cells were stained with both primary and secondary antibodies (e.g. for example, as a negative control for Nanog we stained day 2-3 differentiated cells). For each image, the percentage of positive cells for each channel was calculated. The statistical method is indicated in the figure caption.

### Live bioluminescent imaging

#### Sample preparation

Calibration cells (PGK-Nluc-Fluc - Mandic et al., 2017) and cells carrying Spry4-Fluc gene trap and Nanog::Nluc fusion were routinely cultured in 2i/LIF as described above. Imaging experiments were performed in FluoroDishes (WPI, FD35-100) coated with CellAdhere (now discontinued) for 30min in the incubator before washing gently twice with DPBS with calcium and magnesium (Thermo Scientific/Gibco, 14040133). 100,000 cells/cm^2^ reporter cells and 10,000 calibration cells were plated overnight in 2i medium, before washing gently with DPBS with calcium and magnesium and changing media to N2B27 supplemented with DMSO or RSK(i), 0.5 mM Luciferin (to visualise Fluc, NanoLight Technology, 306A) and 1:2000 RealTime-Glo MT Cell Viability Assay Substrate (to visualise Nluc, first diluted 1:1 in DMSO, Promega G9711).

#### Image acquisition

Bioluminescence live imaging was performed on an Olympus LV200 with an EMCCD camera at 60x (oil-immersion) under environmental control (37°C, 7% CO2) using binning 1x1 (512 x 512) and photon imaging mode 5x. We acquired two images per channel (Fluc: 600 nm LP filter, Nluc: 460/36 nm filter), one with a shorter integration time to measure the bioluminescence signal of calibration cells, one at longer integration time to measure the reporters’ bioluminescent signal. The acquisition times for Fluc were 20ms and 310ms respectively, and for Nluc were 20ms and 240ms.

### Image processing and quantification

Image quantification was performed semi-automatically. The background was subtracted for each image to remove optical aberrations. Next, we generated a maximum intensity mask merging both Fluc and Nluc channels to identify objects. We used a custom Fiji plugin to identify regions of interest (ROIs, 4-pixel diameter circles) in the maximum intensity mask before measuring the intensity in the Nluc and Fluc channel. Cell division was annotated manually. We also measured the background around each ROI for each frame (empty field 10-pixel diameter circle next to the cell ROI).

### Signal normalisation

Since Nluc signal intensity increases over time because of the progressive increase in available substrate in cells (Mandic et al., 2017) for each experiment we included a 1:10 dilution of calibration cells. Calibration cells constitutively express a NLuc-Fluc fusion construct (Mandic et al., 2017). For each slide, we calculated the normalisation ratio by dividing the Fluc over the Nluc signal for the calibration cells before applying a Loess regression. Next, we applied a rolling average to smooth the background signal. The Fluc reporter signal was normalised by subtracting the raw signal to the smooth background, and denoised by fitting a smoothing spline regression. The Nluc signal was normalised by subtracting the raw Nluc signal to the background and dividing by the normalisation ratio and denoised using a low pass filter.

### Reconstructing promoter activity

We used a previously established approach to infer promoter activity from the de-noised Spry4 reporter readout (Kannan et al., 2018). Briefly, this method relies on a series of ordinary differential equations to infer the promoter activity given a resulting output (in this case bioluminescence signal) and a set of fixed parameters. The key parameters are the Fluc degradation rate (d, inferred from data = 0.6), the translation rate (beta, estimated from the average translation rate in ESCs (Ingolia et al., 2011) = 5.59) and the mRNA degradation rate (d.r, average= 0.075). We assume that the magnitude of the maturation rate of bioluminescent proteins is negligible.

### Autocorrelation between Spry4 promoter activity and Nanog downregulation

First, we filtered the dataset to exclude timepoints when Nanog is already downregulated. This removed bias between fast and slow differentiating cells. To obtain a combined dataset, we joined the traces of each cell end-to-end. Next, we generated two control datasets. For the randomised control dataset, the Fluc promoter signal was completely randomised for each cell. For the time shifted control dataset, a random frame shift to was added to the Fluc promoter signal. Next, we calculated the autocorrelation function (ACF) between the smoothened Nanog::Nluc signal and the smoothened Fluc signal as well as the randomised and time-shifted Fluc controls. To determine if the was a significant correlation, we compared the maximum ACF for each trace of the data against the randomised control (noise) and performed a Wilcoxon test. To determine if there was a consistent and significant lag (time relationship) between the Fluc promoter signal and Nanog downregulation, we compared the lag of the curves which show significant autocorrelation (above noise) against the time shifted controls (where the time relationship between the two signals should be random). We compared the distributions using a Kolmogorov–Smirnov test.

### CUT&Tag

#### Sample preparation

Each batch consists of twelve samples processed in parallel. In each batch, the three samples [2i, N2B27/DMSO+siControl, N2B27/MEK(i)+siNanog] were processed in parallel for a given antibody, as well as 2 negative controls and one positive control (typically H3K27me3). Each sample consists of a single well of a 12-well plate. Cells were differentiated as described above and processed following the benchtop CUT&Tag protocol v2 (Kaya-Okur et al., 2019) dx.doi.org/10.17504/protocols.io.z6hf9b6) with minor modifications: all eppendorfs were pre-coated with PBS +0.1% BSA for 30min and all buffers were kept on ice prior to use. Nextera primers were used to generate libraries following the benchtop CUT&Tag recommended protocol. Antibodies and primers used are listed in Table S1. Libraries were pooled and sequenced 50bp pair-end Nextera.

### Alignment and normalisation

Sample were processed in Galaxy (Afgan et al., 2022) workflows available in https://github.com/CMulas/CUT-Tag_analysis). FASTA files were aligned using Bowtie2 (Langmead and Salzberg, 2012) against mouse (mm10) and Escherichia coli K12 (spike in control). BAM files were then sorted, removing unaligned reads and PCR duplicates. For each antibody, all samples were normalised to 1x using bamCoverage (deepTools, (Ramírez et al., 2016).

### Differential binding analysis

For each sample, peaks were called using SEACR (Meers et al., 2019). To determine differential binding for a given antibody we first generated a file of all possible binding regions across samples. To generate this list, all individual lists of peaks were concatenated, sorted and merged. Next, we centred the peaks and trimmed all regions to 1000bp. Finally, we generate a count matrix of the total signal over the 1000 bp regions for each sample. Differential analysis was performed using EdgeR (Robinson et al., 2010). For H3K27me3, we divided regions into promoters and enhancers based on overlap with transcriptional start site (TSS). Peaks were annotated using PAVIS (Huang et al., 2013). The code and list of differentially bound sites can be found in https://github.com/CMulas/2022-Mulas_NanogERK.git.

## Acknowledgements

We thank Peter Humphries and Darren Clements for assistance with microscopy, Brian Hendrich, Nichola Reynolds, Masaki Kinoshita, Jenny Nichols and Lawrence Bates for discussions. Elena Corujo-Simon provided TS cells for positive control RT-qPCR. Pre-conjugated pA-Tn5 was a kind gift from the Henikoff lab. We also thank the Genomics and Tissue Culture facility at the Wellcome-MRC Cambridge Stem Cell Institute for support. CM was funded by the King’s Prize Fellowship at the Randall Centre for Cell and Molecular Biology, King’s College London and a travel grant by British Society of Developmental Biology/The Company of Biologist. AS is a Medical Research Council Professor (G1100526/2). This work was also funded by a Leverhulme Trust Grant (RPG-2016-418, AS and KC) and by European Research Council grant ‘CellFateTech’, 772798 (to KC). The Wellcome–MRC Cambridge Stem Cell Institute receives core support from Wellcome and the Medical Research Council.

---

**Figure S1.**
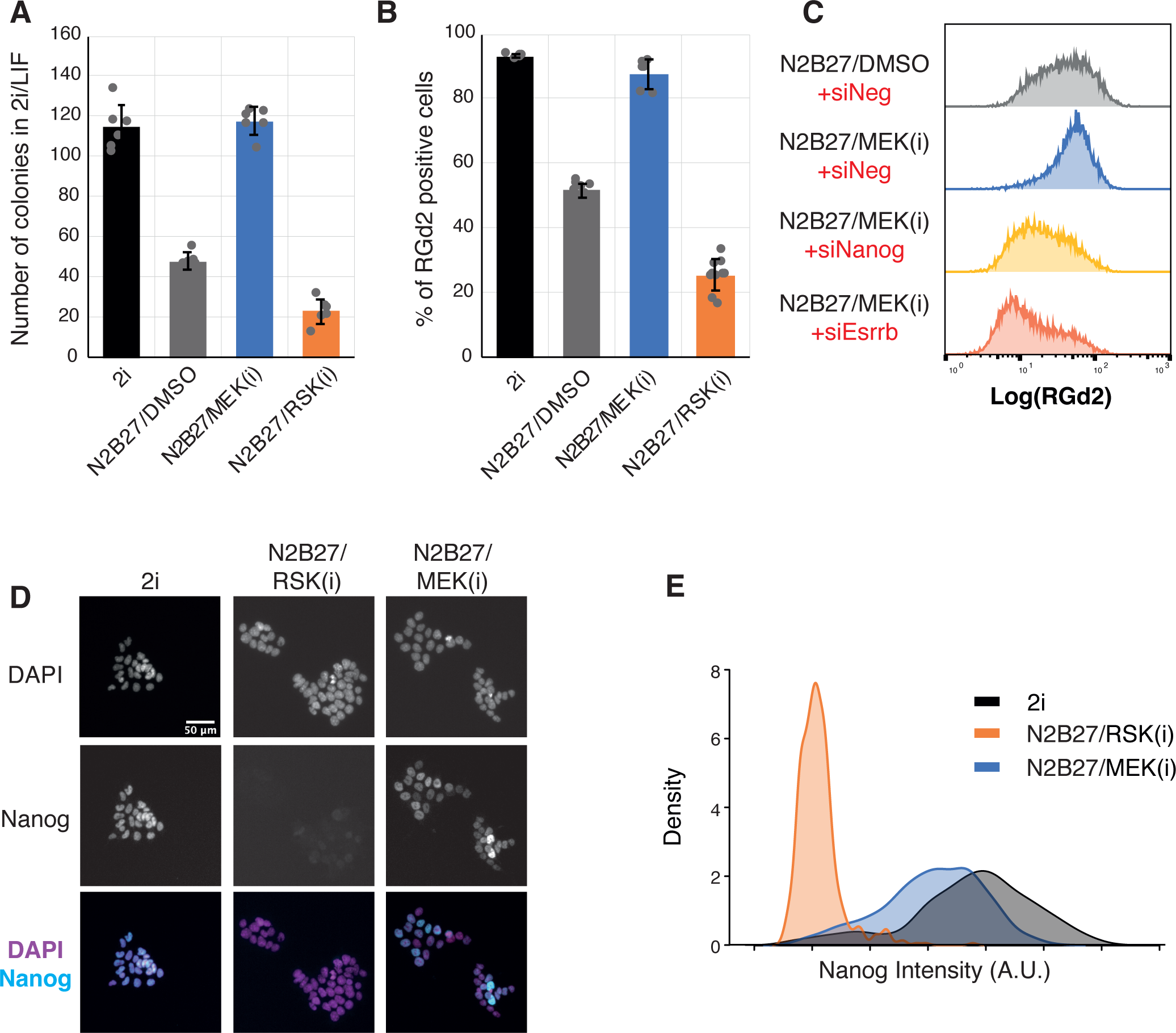
(A) Clonal assay after 30hrs in N2B27 supplemented with 2i, DMSO (control), MEK(i) or RSK(i). Mean ± SD (N=3). (B) Percentage of RGd2 positive cells at 30hrs in N2B27 supplemented with 2i, DMSO (control), MEK(i) or RSK(i). Mean ± SD (N=8). (C) Representative distributions of RGd2 expression from flow cytometry analysis under different conditions. (D) Representative images of immunofluorescence for Nanog at 22hrs in different conditions. Nuclei were counterstained with DAPI. (E) Density distribution plots of mean nuclear Nanog intensity at 22hrs in different culture conditions.

**Figure S2.**
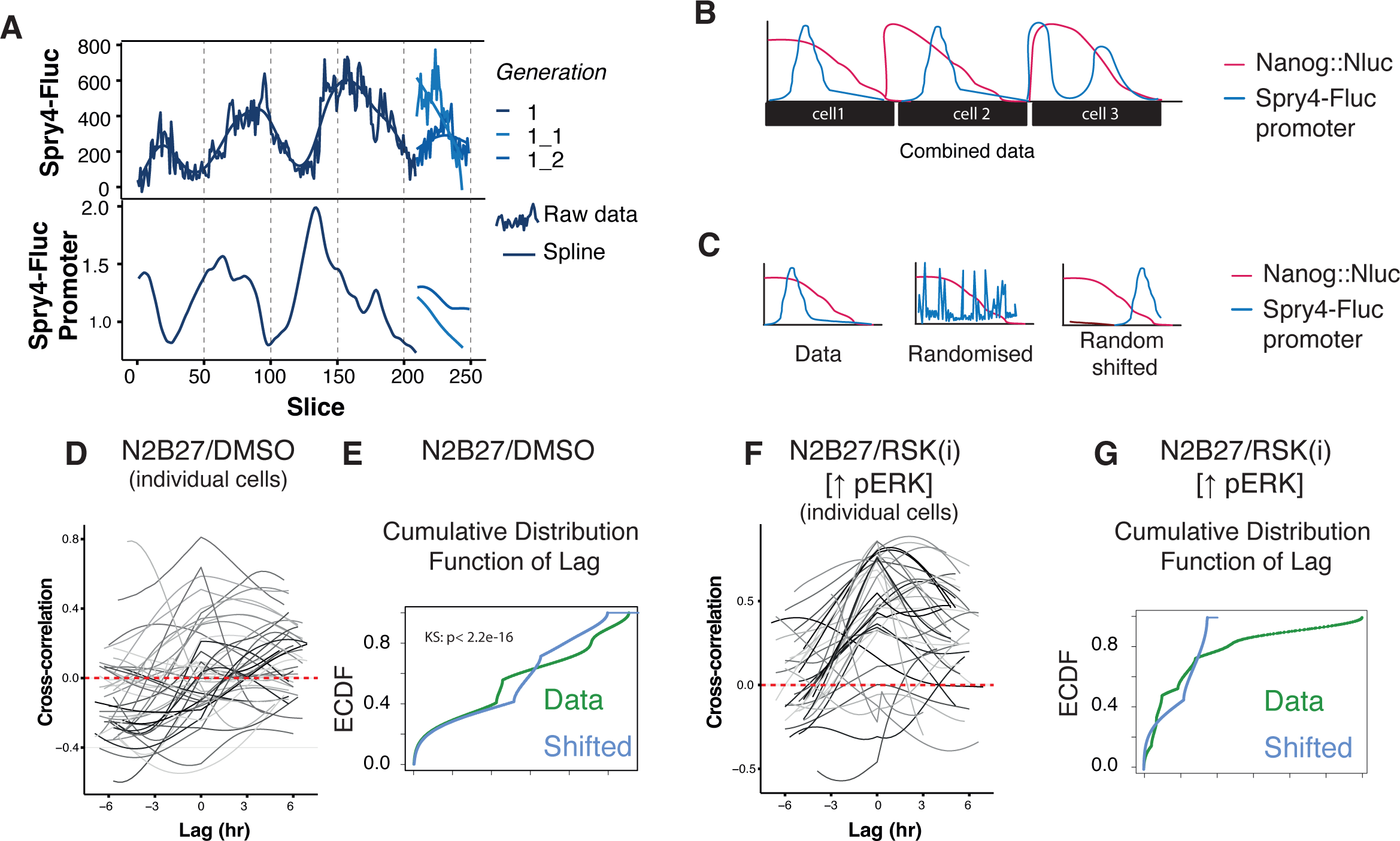
(A) Representative example of Spry4 raw data, spline fit through the raw data (top panels) and calculated Spry4-Fluc promoter activity. (B) Schematic of the combined dataset used to calculate overall autocorrelation function between Spry4-Fluc promoter activity and Nanog protein. (C) Schematic of the controls used. For the randomised control, the Spry4-Fluc promoter signal was completely scrambled. For the random shifted control, to each cell we applied a random shift to the Spry4-Fluc promoter signal. (D) Autocorrelation plot of individual cells in N2B27/DMSO conditions. (E) Cumulative distribution function of the peak lag for cells that show a significant ACF above noise in N2B27/DMSO conditions. (F) Autocorrelation plot of individual cells in N2B27/RSK(i) conditions. (G) Cumulative distribution function of the peak lag for cells that show a significant ACF above noise in N2B27/RSK(i) conditions.

**Figure S3.**
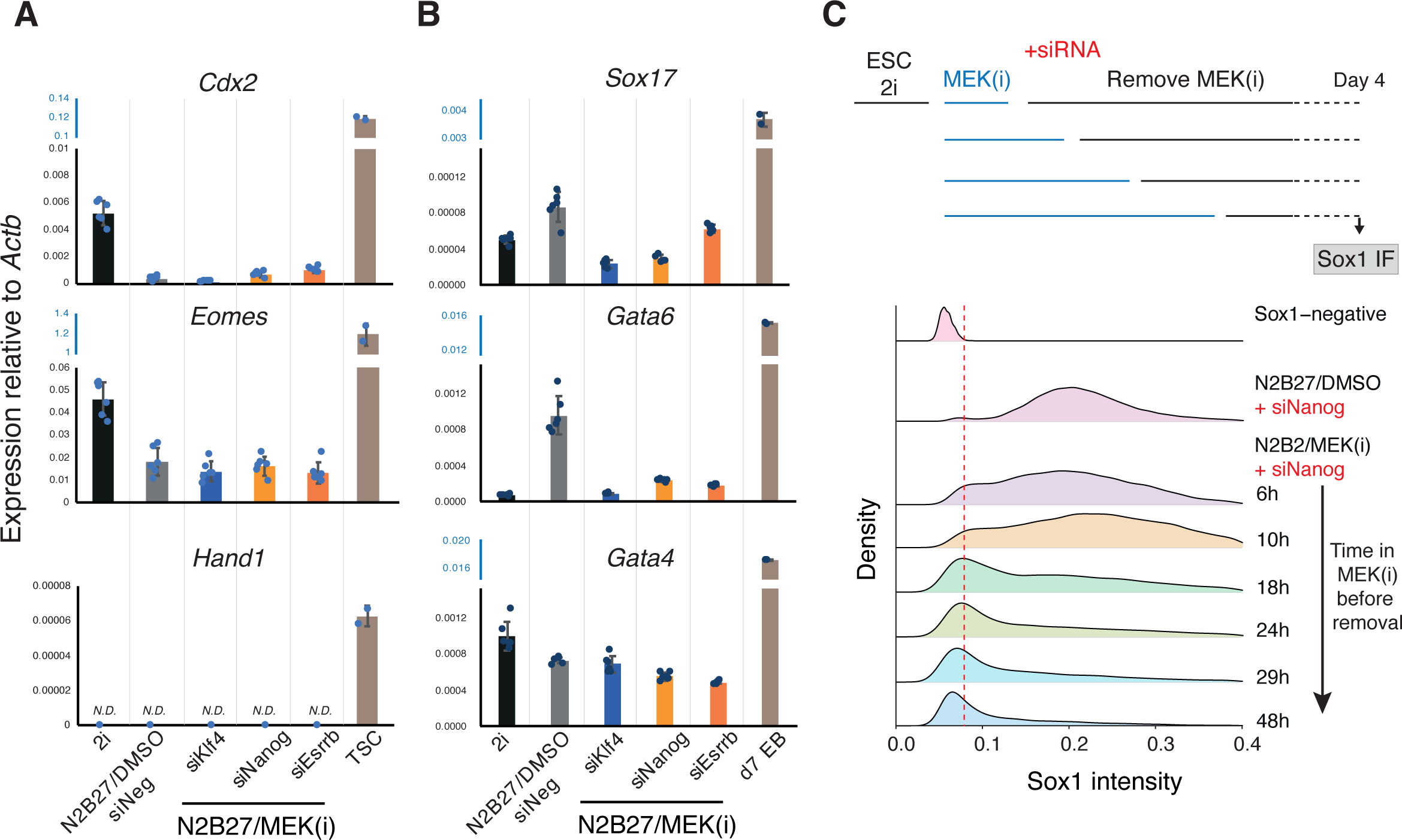
(A) Expression of trophectoderm-associated genes. Expression normalised to *Actb* compared to trophectoderm stem cells (TSC). N.D.: not detected. Mean ± SD (N=4). (B) Expression of primitive endoderm-associated genes. Expression normalised to *Actb* compared to day 7 embryoid bodies. N.D.: not detected. Mean ± SD (N=4). (C) Neural differentiation of siNanog cells following increasing length of MEK(i) treatment. Representative density plots showing distribution of mean Sox1 nuclear intensity on day 4.

**Figure S4.**
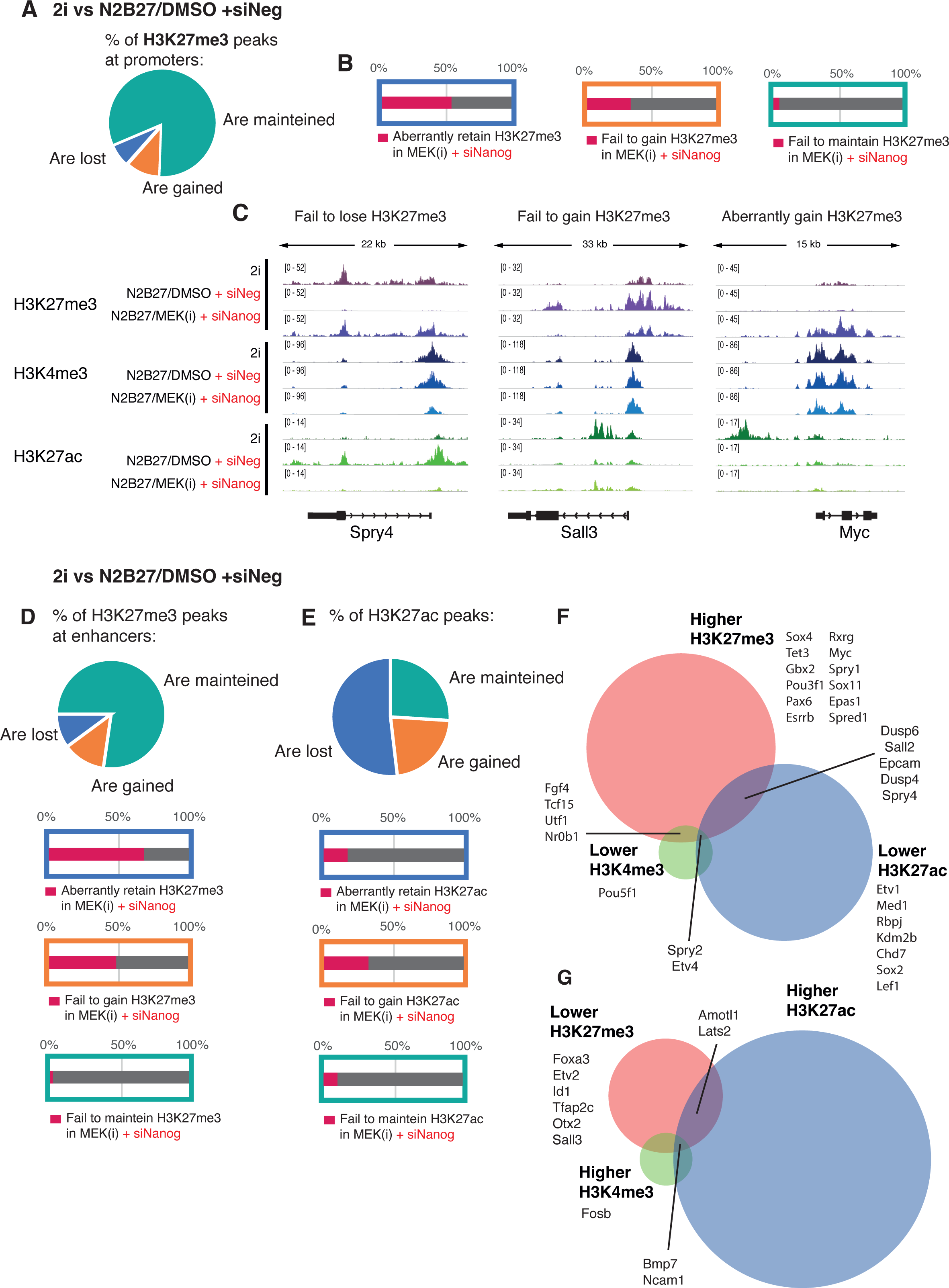
(A) Percentage of H3K27me3 promoter peaks that change (FDR < 0.01, |logFC| > 1) or are maintained during normal differentiation (2i vs N2B27/DMSO + siNeg). (B) Proportion of peaks that show a significant difference during normal differentiation vs N2B27/MEK(i) + siNanog cells. (C) Distribution of H3K27me3, H3K4me3 and H3K27ac at representative loci that fail to follow the expected patterns observed in control conditions. (D) Top panel: Percentage of H3K27me3 enhancers peaks that change (FDR < 0.01, |logFC| > 1) or are maintained during normal differentiation (2i vs N2B27/DMSO + siNeg). Bottom panel: proportion of peaks that show a significant difference during normal differentiation vs N2B27/MEK(i) + siNanog cells. (E) Top panel: Percentage of H3K27ac enhancers peaks that change (FDR < 0.01, |logFC| > 1) or are maintained during normal differentiation (2i vs N2B27/DMSO + siNeg). Bottom panel: proportion of peaks that show a significant difference during normal differentiation vs N2B27/MEK(i) + siNanog cells. (F) Regions likely to be repressed in N2B27/MEK(i) + siNanog cells. Overlap in the regions that show significantly (FDR < 0.01, |logFC| > 1) higher H3K27me3, lower H3K27ac or lower H3K4me3. (G) Regions likely to be activated in N2B27/MEK(i) + siNanog cells. Overlap in the regions that show significantly (FDR < 0.01, |logFC| > 1) lower H3K27me3, higher H3K27ac or higher H3K4me3.

**Figure S5.**
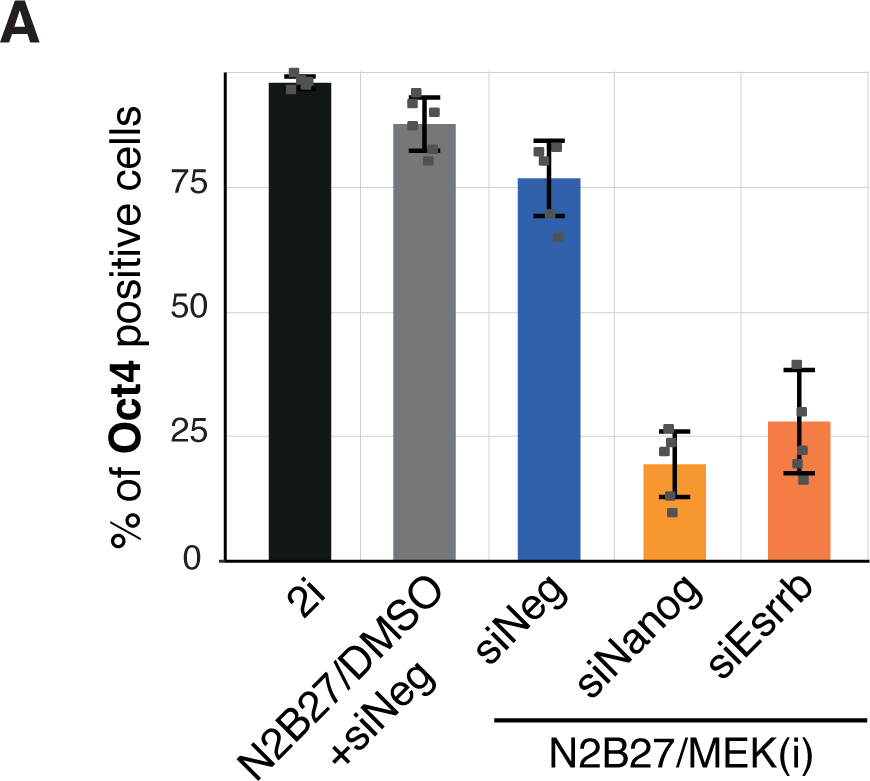
(A) Percentage of Oct4 positive cells at 30hrs under different culture conditions.

**Table S1.** List of antibodies and primers.

